# A comparative study of two grasswren species reveals strong genetic divergence between a peninsula and mainland population

**DOI:** 10.1101/2022.08.15.503954

**Authors:** Aline Gibson Vega, Michelle L. Hall, Amanda Ridley, Saul J. Cowen, Amy L. Slender, Allan H. Burbidge, Marina Louter, W. Jason Kennington

**Affiliations:** School of Biological Sciences, The University of Western Australia, Crawley, WA 6009, Australia; Bush Heritage Australia, 1/395 Collins St, Melbourne, Victoria, Australia; Biodiversity and Conservation Science, Department of Biodiversity, Conservation and Attractions, Locked Bag 104, Bentley Delivery Centre, WA 6983, Australia; College of Science and Engineering, Flinders University, Adelaide, SA, Australia; School of Science, Edith Cowan University, 100 Joondalup Drive, WA 6027, Australia

## Abstract

Dispersal patterns dictate genetic population structure, and ultimately population resilience, through maintaining critical ecological processes and genetic diversity. Direct observation of dispersal events is not often possible, but genetic methods offer an alternative method of indirectly measuring dispersal. Here, we use 7 652 genome-wide single-nucleotide polymorphisms (SNPs) to evaluate genetic population structure and infer dispersal capabilities of the Western Grasswren *(Amytornis textilis textilis;* WGW) in Western Australia (*n* = 118), utilising a sister species, the Thick-billed Grasswren *(Amytornis modestus;* TBGW) as a comparison dataset (*n* = 80). We found genetic divergence and low genetic diversity between two populations (Hamelin and Peron) in the WGW, despite evidence of long dispersal distances within populations by females. In addition, the two WGW populations were found to be more genetically divergent than two described subspecies of TBGW, despite the WGW occurring over a smaller spatial scale. By comparing these two grasswren species, our data suggest a narrow strip of land may be acting as a geographic barrier in the WGW, limiting dispersal between a peninsula population to the mainland. We investigate if morphology aligns with genetic divergence, with some estimates of divergence between WGW populations greater than those between subspecies of TBGW. However, confidence intervals were large, preventing definitive conclusions. Our results support the hypothesis that peninsula populations of small, ground-dwelling birds are genetically isolated from adjacent mainland populations. Furthermore, there is evidence to suggest that the limited gene flow is asymmetrical, with directional dispersal occurring from the bounded peninsula population to the mainland. Our study also highlights how substantial genetic divergence does not necessarily coincide with phenotypic differences.

## Introduction

Dispersal influences genetic population structure, and hence population resilience through maintaining genetic diversity and buffering against genetic drift (Frankham 1996; Frankham *et al*. 2002). Dispersal itself can be influenced by biotic and abiotic factors. Some species tend to exhibit short-distance dispersal behaviours, causing strong genetic population structure in an otherwise homogenous landscape (Aguillon *et al*. 2017). Geographic features may also restrict an individual’s capacity to disperse. Landscape features such as highly heterogeneous or fragmented habitat (i.e. islands, peninsulas, roads, artificial waterways and habitat clearing) can be geographic barriers that restrict gene flow over relatively short spatial scales. For example, populations of Dibbler *(Parantechinus apicalis),* a small marsupial, separated by small cliffs and unsuitable vegetation have been found to be genetically subdivided over relatively short spatial scales (19 km) (Thavornkanlapachai *et al*. 2019a). In an arboreal species, a barrier to dispersal was found in the Western Ringtail Possum *(Pseudocheirus occidentalis),* where a narrow artificial waterway (30 m wide) was found to be a greater barrier than a major road an equal distance away (200 m) (Yokochi *et al*. 2016). Even in small passerines, surprising barriers to dispersal can be found. In White-fronted Chats *(Epthianura albifrons),* unsuitable habitat was found to be less of a barrier to gene flow than urbanisation, despite the urban populations being substantially geographically closer (20 km) than those separated by unsuitable continuous forest (500 km) (Major *et al*. 2014). These few examples highlight how some landscape features can present themselves as greater barriers to dispersal than others, and that it is very much species-specific.

Shark Bay is a semi-arid region in Western Australia with a variety of landscape features such as dune systems and variable habitat (Payne *et al*. 1987). The presence of numerous narrow inlets and broad gulfs creates a complex coastline throughout the region. The terrestrial parts of Shark Bay encompass a peninsula (Peron Peninsula) connected to mainland Shark Bay via an *Acacia* sandplain isthmus (Brooker 2000; Figure 1). The isthmus measures less than 2.5 km at the narrowest point. The narrow terrestrial connection of Peron Peninsula and complex coastline separating it from the mainland are thought to be a potential dispersal barrier for some small passerines, such as the White-winged Fairywren (*Malurus leucopterus;* Walsh *et al*. 2021), as limited habitat is present across the isthmus and dispersal across open water is unlikely for small passerines. In that broad-scale study, there was a lack of obvious genetic differentiation between mainland White-winged Fairywren populations sampled almost 2500 km apart, yet samples from Peron Peninsula and the Shark Bay mainland sampled only 100 km apart showed significant genetic differentiation, suggesting that there may be a barrier to dispersal between the peninsula and mainland (Walsh *et al*. 2021). Yet, it remains unclear whether other species in this region exhibit similar patterns.

**Figure 1.**
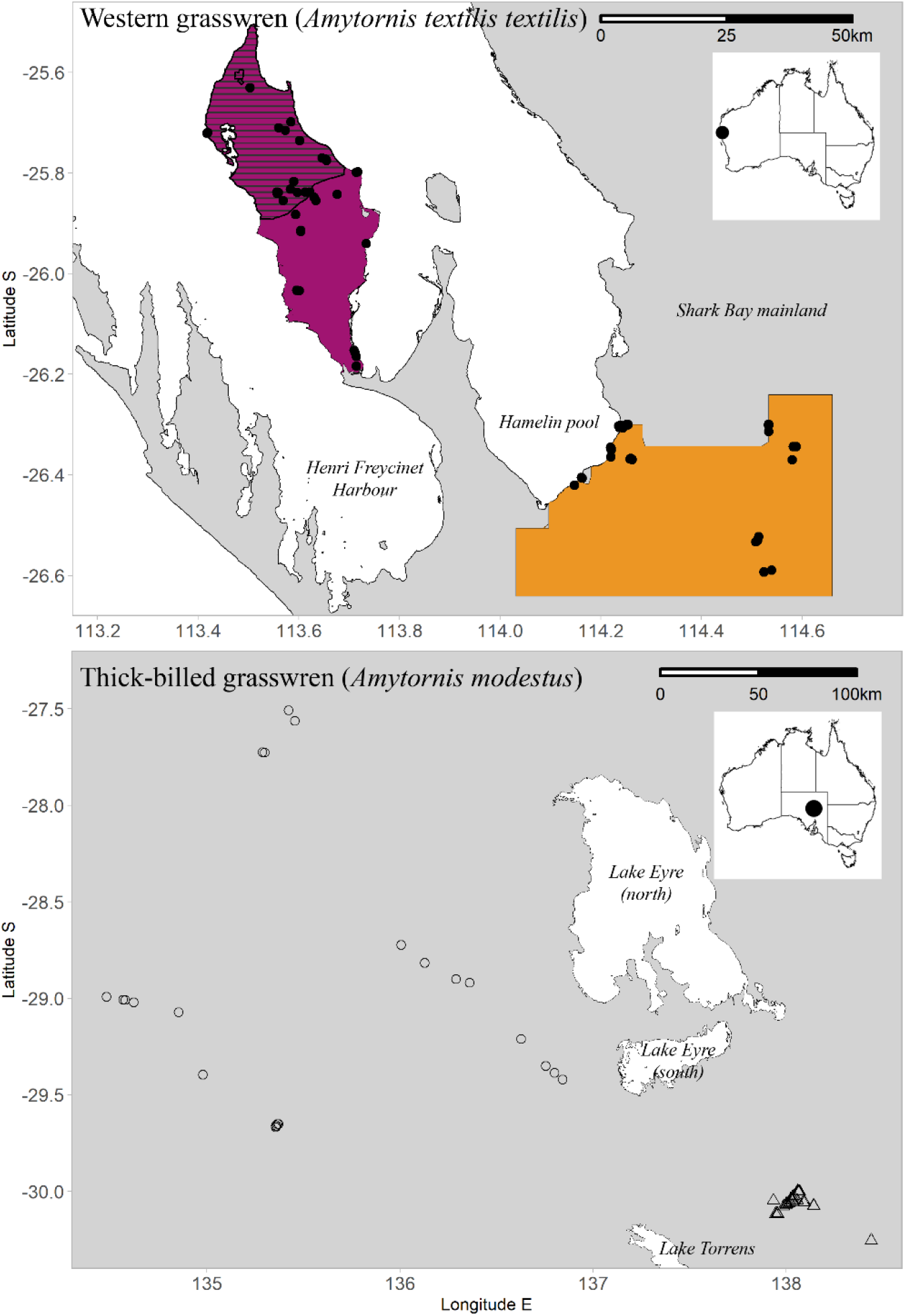
Location of western grasswren (WGW; *Amytornis textilis textilis)* and thick-billed grasswren (TBGW; *Amytornis modestus)* samples. In the WGW map, purple and orange shaded areas indicate areas of interest where blood samples were attempted to be collected from (Peron Peninsula and Hamelin Station Reserve, respectively), with the hatched area in Peron Peninsula denoting Francois Peron National Park. Closed circles indicate collection locations of blood samples from *Amytornis textilis textilis.* In the TBGW map, open circles and open triangles indicate original collection location of blood samples from *Amytornis modestus indulkanna* and *A. m. raglessi,* respectively. The white in the WGW map denotes ocean while in the TBGW map it denotes Lake Eyre and Lake Torrens.

Here, we investigate population structure of a small ground-dwelling passerine, the Western Grasswren, subspecies *Amytornis textilis textilis* (WGW), in the Shark Bay, region. This subspecies occupies two disjunct areas in Shark Bay, Peron Peninsula and the adjacent mainland near Hamelin Pool (Figure 1). While there has been considerable focus on resolving and redefining the taxonomy of *Amytornis* (e.g. Black *et al*. 2010; Christidis *et al*. 2013), little is known about grasswren ecology, with only one grasswren species having been studied in depth (Thick-billed Grasswren, *Amytornis modesuts;* Louter 2016; Slender 2018). Grasswren dispersal movement and consequent population structuring is not well understood. Based on the strong genetic structuring between White-winged Fairywren populations in Peron Peninsula and Hamelin Pool (Walsh *et al*. 2021), a passerine with morphological and behavioural similarities to grasswrens, we hypothesise that WGW populations in these areas will also show strong genetic structuring. To provide context for the genetic data, we compared the two Western Grasswren populations in Western Australia with two subspecies of a sister species in South Australia, the Thick-billed Grasswren (TBGW; *Amytornis modestus indulkanna* and *A. m. raglessi*). It should be noted that the TBGW has a different evolutionary history to the WGW, with the TBGW being a parapatric species having made secondary geographical contact in evolutionary recent times (Slender 2018). The two subspecies have also been argued to belong to two separate evolutionary significant units (ESUs) (Austin *et al*. 2013), however its current lack of genetic differentiation and high level of gene flow has led to the suggestion that they could be regarded as one ESU for conservation purposes (Slender *et al*. 2021). In contrast, the WGW populations remain as one ESU and have likely been in geographical contact throughout much of their evolutionary history (Payne *et al*. 1987; Austin *et al*. 2013).

While Thick-billed Grasswren habitat is slightly different to habitat found in Shark Bay, the key habitat requirements are similar for both Western and Thick-billed Grasswrens (Brooker, 2000; Slender, 2018). Both Western and Thick-billed Grasswren species require low shrub vegetation such as *Acacia* and Chenopod shrub land, respectively, which provide low-to-the-ground cover for foraging, shelter and nesting (Brooker 1998; Louter 2016). The distribution of both grasswren species encompass areas of variable suitable plant communities mixed with patchy unsuitable habitat; however, the WGW landscape has added complexity, with coastline being a prominent feature.

In this study, we use genome-wide single nucleotide polymorphisms (SNPs) to evaluate population genetic structure, infer dispersal capabilities and identify any landscape barriers in the WGW and TBGW. First, we aim to quantify genetic divergence between the two species to confirm that the TBGW represents an independent comparison dataset to the WGW, consistent with previous mtDNA studies. Second, we describe how genetic variation is partitioned within and between populations within each species and assess evidence of admixture between populations within each species. Third, we explore fine-scale dispersal patterns within each species and test for evidence of sex biased dispersal, which is a common phenomenon in many taxonomic groups (Greenwood 1980). Last, we looked for evidence of morphological divergence between populations within each species, predicting again that divergences will be greater in WGW due to greater landscape complexity causing greater isolation between populations.

## Materials and methods

### Study species

The Western Grasswren *(Amytornis textilis)* is a ground-dwelling arid zone passerine, which occurs around Shark Bay, Western Australia, and within and around the Gawler Ranges in South Australia (Brooker 2000; Black *et al*. 2009). This study focuses on the Shark Bay subspecies, *A. textilis textilis* (WGW). Under the Western Australian *Biodiversity Conservation Act 2016,* the Western Australian subspecies *A. t. textilis* is listed as a Priority 4 species, defined as ‘rare, near threatened and in need of monitoring’. Two large conservation properties are strongholds for the two remaining grasswren populations in Shark Bay. Francois Peron National Park extends over half of Peron Peninsula (Peron) and is managed by the Department of Biodiversity, Conservation and Attractions, and Hamelin Station Reserve (Hamelin), which is owned and managed by Bush Heritage Australia as a conservation reserve, is in the broader mainland landscape south-east of the peninsula (Figure 1).

The second species examined, the Thick-billed Grasswren *(Amytornis modestus)* is morphologically, genetically and geographically distinct from WGW (Black *et al*. 2010; Austin *et al*. 2013). The TBGW is listed as ‘vulnerable’ under the Commonwealth *Environment Protection and Biodiversity Conservation Act 1999.* The two subspecies of interest within the TBGW, *A. m. indulkanna* and *A. m. raglessi* (hereafter referred to as Indulkanna and Raglessi respectively and collectively as TBGW), are distinguishable based on their morphological and mitochondrial differences (Black *et al*. 2010; Austin *et al*. 2013). Indulkanna is found west of Lake Eyre and Lake Torrens, South Australia, while Raglessi can be found on the periphery of the northern Flinders Ranges (Black 2011; Figure 1).

### Sampling

Western Grasswrens were captured in Shark Bay in 2019 and 2020 using mist-nets. Blood samples were collected from the brachial vein and stored in 100% ethanol. Samples were obtained throughout the peninsula and on parts of mainland within the known WGW distribution (Figure 1). A total of 63 and 55 samples from adult individuals were obtained from Peron and Hamelin respectively. Juvenile grasswrens, identified by morphological features in-hand, were not included in the dataset as they have not had the opportunity to disperse, and thus would bias the data.

DNA samples of Indulkanna and Raglessi were sourced from the South Australian Museum. These DNA samples were originally sourced from blood tissue stored in 100% ethanol. The collection and DNA extraction process for these samples is described in Slender *et al*. (2017a). A total of 47 Raglessi and 33 Indulkanna samples with known geographic location were used in this study (Figure 1).

### SNP genotyping

Blood and DNA samples were sent to Diversity Arrays Technology for genome-wide SNP sequencing using a genotype-by-sequencing approach (DArTseq; http://www.diversityarrays.com/). This approach involves a combination of DArT complexity reduction methods and next-generation sequencing platforms (Kilian *et al*. 2012). Several enzyme systems for complexity reduction were tested and the *Pstl-Sphl* method was chosen for *Amytornis.* The *Pstl-*compatible adaptor was comprised of an Illumina flowcell attachment sequence, a sequencing primer and a staggered barcode region of varying lengths. The reverse adapter (*Sphl-*compatible) contained the Illumina flowcell attachment sequence and a *Sphl* overhang sequence. Only fragments with both *Pstl* and *Sphl* were amplified by polymerase chain reaction (PCR) with an initial denaturing step at 94 °C for 1 min, followed by 30 cycles of temperature changes as follows: denaturation at 94 °C for 20 seconds, annealing at 58°C for 30 seconds, and extension at 72 °C for 45 seconds, with an additional final extension at 72 °C for 7 min (Melville *et al*. 2017).

Following PCR amplification, the products from each sample were bulked and applied to c-Bot (Illumina) bridge PCR followed by sequencing on Illumina Hiseq2500. The sequencing (single read) was run for 77 cycles. Sequences were processed using proprietary DArT analytical pipelines. In the primary pipeline the ‘fastq*’* files were first processed to filter away poor quality sequences, applying more stringent selection criteria to the barcode region compared to the rest of the sequence. Identical sequences were collapsed into ‘fastqcoll files’ and groomed using DArT’s proprietary algorithm to correct low quality bases from singleton reads, using collapsed tags with multiple members as a template. The ‘groomed’ fastqcoll files were used in the secondary pipeline for SNP calling (Kilian *et al*. 2012).

Following the generation of 134,726 SNP loci, multiple SNP loci on the same contig, loci that were genotyped in fewer than 95% of samples, had a minor allele frequency less than 0.05, or were monomorphic were removed prior to analysis. Potential outlier loci were screened using three detection methods: BAYESCAN 2.1 using default settings (Foll and Gaggiotti 2008), outflank (Whitlock and Lotterhos 2015) and PCADAPT (Luu *et al*. 2017). Multiple detection methods were utilised as each method has a different threshold for identifying outliers. The outlier screening was done independently for WGW and TBGW. Outlier loci were identified by having q-values less than 0.05 and having been detected across at least two outlier detection methods. These loci were removed from the global dataset prior to assessments of population structure and genetic diversity. Relatedness was determined through GenAlEx 6.503 (Peakall and Smouse 2006; Peakall and Smouse 2012) using the Ritland (1996) *r* estimate. One individual from a pair having a relatedness coefficient of 0.2 or more was removed for each species’ respective genetic analysis to exclude highly related individuals from subsequent analyses. A sensitivity analysis for the filtered loci was conducted using more conservative filtering parameters (see Appendix S1 in Supporting Information).

### Morphology

Morphological divergence within the WGW population in the Shark Bay area has not previously been examined. While we could have tested for direct differences between Hamelin and Peron, we were specifically interested in comparing the magnitude of divergence within the WGW to that in the TBGW, utilising the TBGW as a ‘benchmark’ for what constitutes significant morphological divergence for subspeciation. Morphological measurements were obtained for all grasswrens at their time of capture. Head-bill (back of head to tip of bill) and tarsus (intertarsal joint to base of toes) were recorded using dial callipers (± 0.05 mm). Wing length (carpal joint to the tip of the longest primaries, flattened against the ruler) and tail length (base to tip of the longest tail feather) were recorded using a ruler (± 0.1 mm). As grasswrens spend a lot of time in the undergrowth of shrubs, tail feathers can often be shortened due to wear and damage; measurements of damaged tails, which had obviously lost length, were not included in the analysis. Most grasswren weights were recorded using a 100 g Pesola spring scale (± 0.5 g), with the exception of all TBGW and 25 WGW individuals, which were weighed on a digital scale (± 0.1 g). Measurements were recorded by multiple people, with A. Gibson Vega measuring most of the WGW (> 50% of the samples). The TBGW were measured by A. L. Slender (Indulkanna) and M. Louter (Raglessi).

### Data analysis

#### Population structure and genetic diversity

Data analyses were performed using R v4.0.3 unless specified (R Core Team 2020). Estimates of observed and expected heterozygosity, inbreeding coefficient (*F*_IS_) and pairwise fixation index (*F*_ST_) were obtained using the R package ‘hierefstat’ (Goudet 2005), following Nei (1987) for the *F*_ST_ values. The significance of *F*_IS_ and *F*_ST_ values were determined by bootstrapping 1000 replicates with a 95% confidence interval. Pairwise *F*_ST_, a measure of population differentiation due to genetic structure, can be misleading in species with low genetic variation. To test for consistency, Hedrick’s *G*’_ST_ was calculated using GenAlEx 6.503 (Peakall and Smouse 2006; Peakall and Smouse 2012), which provides a better metric when comparing loci with varying genetic variation and between organisms with different effective population sizes (Hedrick 2005). *G*’_ST_ significance was determined by bootstrapping 999 replicates with 95% confidence interval. A Mantel test, implemented using the R package ‘ade4’ (Dray and Dufour 2007) was used to test the significance of the correlation between *F*_ST_ and *G*’_ST_ values. Significance of the test was determined by bootstrapping 9999 replicates with a 95% confidence interval. To test for significant differences in observed and expected heterozygosity values between populations, a Friedman test was carried out, where each locus represented a block. A pairwise Wilcoxon rank sum test determined which pairs of populations had significantly different levels of genetic diversity.

The software package STRUCTURE 2.3.4 (Pritchard *et al*. 2000) was used to infer genetic clustering and evidence of admixture. STRUCTURE constructs models using a Bayesian clustering method which estimates the proportion of each individual’s genome having ancestry in each cluster for any given number of clusters (K) (Pritchard *et al*. 2000). This is achieved through Markov-Chain Monte Carlo (MCMC) methods for sampling from a probability distribution. The number of genetic clusters we tested ranged from 1 to 6, with 10 replicates for each value of K. A burn-in of 50 000 followed by 100 000 MCMC replicates was set. We determined the most likely value of K by using the Delta K method (Evanno *et al*. 2005). As this method cannot determine K = 1, we also considered the likelihood probability (Ln (P)), where the most likely K is at the ‘plateau’ of the curve. For these analyses, a hierarchical approach was followed, where all grasswren samples were pooled into the same STRUCTURE analysis. The resulting clustering was then run through STRUCTURE again, but independently for each assigned cluster and controlling for closely related individuals. For the between- and within-species analysis, each individual’s average membership to K clusters from the 10 replicates runs was calculated, re-organised and visualised using the R package ‘pophelper’ (Francis 2017). A sensitivity analysis for the filtered loci was conducted, whereby STRUCTURE analysis for population structure and genetic diversity were re-conducted using more conservative filtering (see Appendix S1).

To test the robustness of the clustering analyses, we also performed a discriminant analysis of principal components (DAPC) using the R package ‘adegenet’ (Jombart *et al*. 2010). DAPC is preferred over the traditional principal component analysis (PCA) as PCA looks for largest overall variance, ignoring prior knowledge on group assignment. In contrast, DAPC maximises variance among groups, while minimising within group variance. This method achieves the best discrimination of individuals into pre-defined groups (Jombart *et al*. 2010). The optimal number of principal components to retain was determined by maximising the highest mean success of successful assignment. The optimal number of clusters was determined using the Bayesian Information Criterion (BIC), where the lowest BIC values are the best model fit.

#### Fine-scale genetic structure

We inferred fine-scale dispersal capabilities by assessing spatial autocorrelation (SA) within each WGW and TBGW population using GenAlEx 6.503 software (Peakall and Smouse 2006; Peakall and Smouse 2012). Non-breeding adults were removed from the WGW dataset, as they have not yet had the opportunity to disperse. Since it was not always known which individuals made up a breeding pair in the TBGW dataset, one female and one male sample were retained per territory. As sample sizes in each distance class were limited, we performed a Distance Class (DC) analysis to calculate and plot mean spatial genetic autocorrelation (*r*) for a series of increasing distance class sizes. The sizes of distance class were variable, depending on the sample size of pairwise geographic distances for each dataset. To test for statistical significance, we used random permutations and bootstrapped *r* using 999 replicates to determine the 95% confidence interval around the null hypothesis (no genetic structure) and error bars, respectively. To investigate sex-biased dispersal the SA analyses were also carried out on each sex separately for the WGW. Low sample sizes precluded single sex analyses for the TBGW.

#### Morphological analysis

Morphometric data were analysed using generalised linear mixed models in the R package ‘lme4’ (Bates *et al*. 2014). A fixed effect of population (Hamelin, Peron, Indulkanna or Raglessi), sex and their interaction on a morphological trait was incorporated, with observer included as a random effect to control for any inter-observer variation. A separate analysis was done for each morphological trait (body mass, head-bill length, tarsus length, tail length and wing length). To test for intraspecific pairwise differences, a post-hoc analysis consisted of a Tukey’s Honest Significant Difference test using the ‘multcomp’ package as it can incorporate mixed models (Hothorn *et al*. 2008). This test was only done on the traits for which a significant effect of population or sex was detected. To compare the magnitude of interpopulation morphological differences between the two grasswren species, pairwise *P*_ST_ values were calculated using the ‘Pstat’ R package for each species (Da Silva and Da Silva 2018). P_ST_ measures the quantitative difference between two groups, similar to *F*_ST_ measures of genetic divergence. 95% confidence intervals were determined by bootstrapping 1 000 replicates through the same package.

## Results

### SNP filtering

Following filtering of the initial 134 726 SNP loci generated by DArTseq, 69 419 loci were removed as they were identified on the same contig as another locus. Other loci that were removed included 44 483 loci with a low call rate, 452 monomorphic loci, 12 710 loci with a low minor allele frequency and 10 outlier loci. The resulting neutral dataset contained 7 652 loci, with 1.84% missing data (no SNP was called for that locus in a particular individual).

### Genetic diversity

Estimates of genetic variation within each population sample are given in Table 1. WGW populations had lower observed (H_o_) than expected heterozygosity (H_e_), as did populations of TBGW (Table 1). Hamelin and Peron had significant differences for *F*_IS_ as well as WGW to TBGW populations (pairwise Wilcox rank sum test; p-value < 0.01) while TBGW had no within-subspecies differences (pairwise Wilcox rank sum test; p-value > 0.05). There was a significant difference in both observed and expected heterozygosity among populations (Friedman’s test; H_o_; *X*^2^ = 679.55, df = 3, p-value < 0.001, He; *X*^2^ = 903.99, df = 3, p-value < 0.001), which a pairwise Wilcox rank sum test indicated were between the two species, with WGW showing lower levels of H_o_ and H_e_ than TBGW, but with no significant differences between the two populations within species (Table 2). *F*_IS_ values were significantly positive in all four populations, indicating heterozygote deficiencies relative to random mating, and were almost two times greater in WGW than in TBGW.

**Table 1.** Summary of descriptive population genetic statistics, observed heterozygosity (Ho), expected heterozygosity (He) and inbreeding coefficient (F_IS_), for the western grasswren *(Amytornis textilis textilis)* and thick-billed grasswren *(Amytornis modestus)* populations. Asterisks indicate significant deviation from zero at p, 0.05 level.

| Species | Population | N | $H_O$ (± SE) | $H_e$ (± SE) | $F_{IS}$ (± SE) |
| --- | --- | --- | --- | --- | --- |
| *A. textilis* | Hamelin | 54 | 0.123 ± 0.002 | 0.142 ± 0.002 | 0.110 ± 0.002* |
| *A. textilis* | Peron | 50 | 0.129 ± 0.002 | 0.145 ± 0.002 | 0.090 ± 0.002* |
| *A. modestus* | Indulkanna | 31 | 0.182 ± 0.002 | 0.232 ± 0.002 | 0.197 ± 0.003* |
| *A. modestus* | Raglessi | 41 | 0.184 ± 0.002 | 0.237 ± 0.002 | 0.201 ± 0.003* |

**Table 2.**
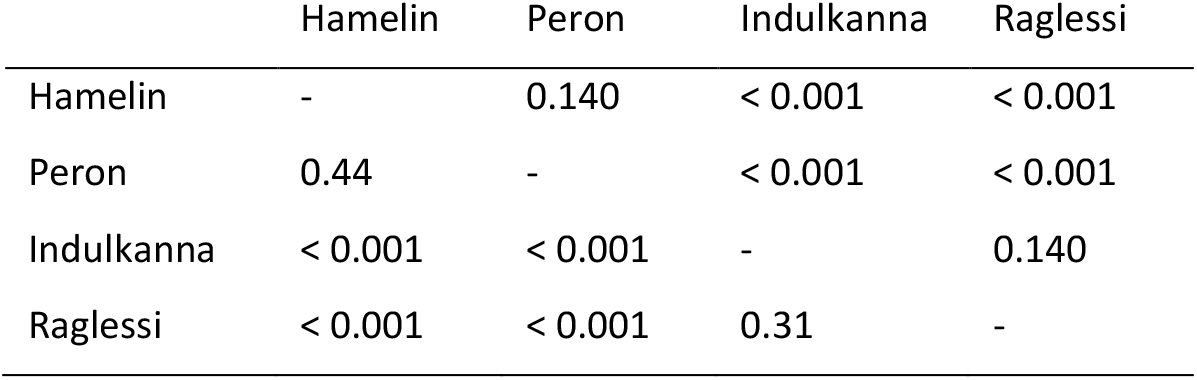
Pairwise Wilcoxon rank sum test results comparing western grasswren *(Amytornis textilis textilis;* Hamelin and Peron) and thick-billed grasswren *(Amytornis modestus;* Indulkanna and Raglessi) populations. Above the diagonal are p values results comparing observed heterozygosity between populations, while below the diagonal p values for expected heterozygosity.

Fourteen pairwise comparisons of WGW individuals (0.002% of 6786 pairwise comparisons) were deemed to be closely related (*r* ≥ 0.2), with one from each pair removed prior to the analysis of population structure. The geographic distance between almost all close relatives ranged from 0 to 1 km. An exception was a pair of individuals, one from Hamelin (male) and one from Peron (female), that were captured 83.5 km (straight line distance) apart. In TBGW, eight pairs had *r* values greater than 0.2, with the greatest distance between pairs being 800 m for Indulkanna and 500 m for Raglessi. As with WGW data, one individual from each pair of close relatives was removed prior to the analysis of population structure.

### Population structure

All four grasswren populations were significantly divergent to each other (Table 3). However, Hamelin and Peron populations showed a greater divergence between each other than Raglessi and Indulkanna populations (Table 3). Pairwise *G*’_ST_ values were highly correlated with the *F*_ST_ values (Mantel test; r = 0.99, P < 0.05), with *G*’_ST_ being more consistently lower than *F*_ST_ in all pairwise comparisons. Significant *G*’_ST_ values were found between WGW and TBGW (*G*’_ST_ = 0.46 – 0.47, *p* = 0.002) and greater interpopulation values between Hamelin and Peron populations (*G*’_ST_ = 0.052, *p* = 0.002) compared to Raglessi and Indulkanna populations (*G*’_ST_ = 0.014, *p* = 0.002). A STRUCTURE analysis including all individual samples showed strong evidence for two genetic clusters (WGW and TBGW), with all individuals within each species assigned to the same genetic cluster and each species assigned to a different genetic cluster. The Delta K analysis also strongly supported the presence of two genetic clusters (Figure 2). The DAPC based on the same individuals also clearly separated the two species (Figure 3), with DAPC BIC identifying the optimal clustering between two and three genetic clusters (see Appendix S2). Interspecific differences heavily loaded towards component one which separated Raglessi and Indulkanna and the grasswren species (PC1; eigenvalue 309.18) while component two loaded towards intraspecific differences in Peron and Hamelin (PC2; eigenvalue 14.62). These two components explain 44.9% of observed variance across all samples.

**Figure 2.**
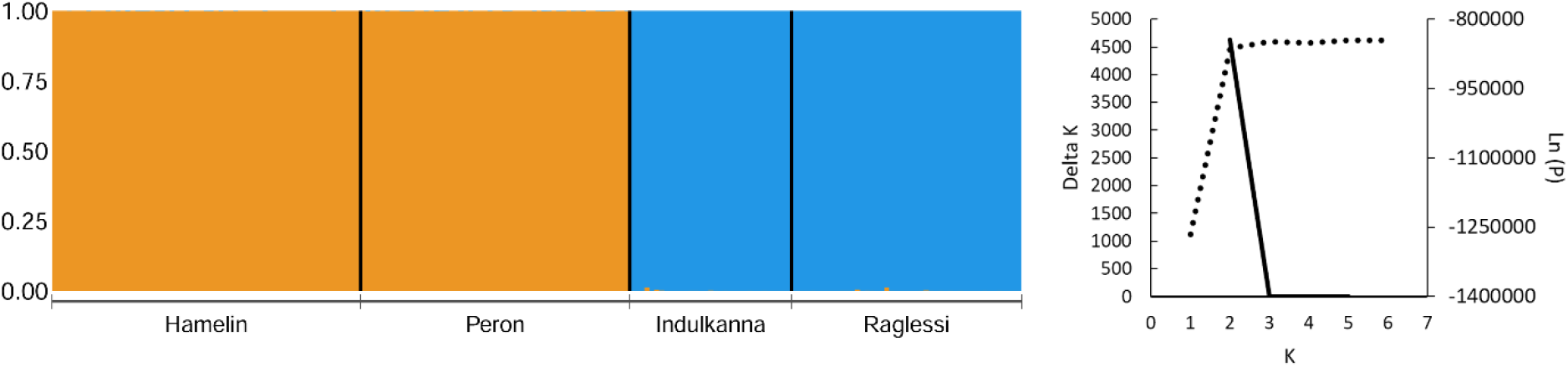
Summary of the STRUCTURE analysis of the western grasswren *(Amytornis textilis textilis;* Hamelin and Peron) and thick-billed grasswren *(Amytornis modestus;* Indulkanna and Raglessi), with genetic clusters represented by different colours (left). Each column represents an individual’s estimated allocation to each cluster. Delta K estimates for varying values of genetic clusters (K; solid line) and probably estimates for each K (Ln (P); dotted line), derived from the STRUCTURE analysis (right).

**Figure 3.**
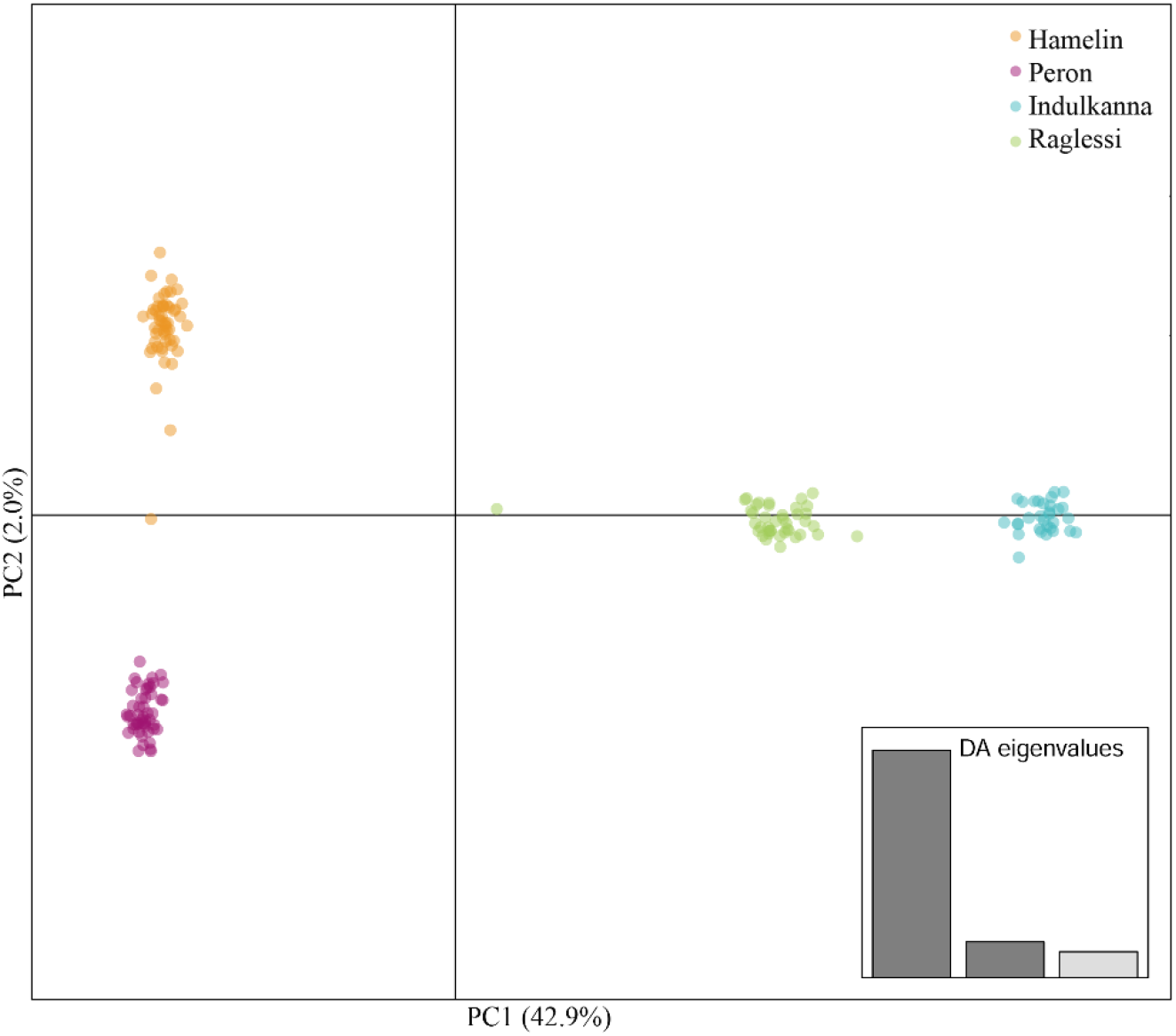
Summary of the discriminant analysis of principal components on the western grasswren *(Amytornis textilis textilis;* Hamelin and Peron) and the thick-billed grasswren *(Amytornis modestus;* Indulkanna and Raglessi), with each population represented by different colours. Proportion of conserved variance for the first two principal components is given in parentheses.

**Table 3.** Pairwise *F*_ST_ (Nei 1987) between populations of western grasswren *(Amytornis textilis;* Hamelin and Peron) and thick-billed grasswren *(Amytornis modestus;* Indulkanna and Raglessi). Below the diagonal are the *F*_ST_ values. Above the diagonal are 95% confidence interval bootstrap results.

|  | Hamelin | Peron | Indulkanna | Raglessi |
| --- | --- | --- | --- | --- |
| Hamelin | - | 0.067 – 0.077 | 0.491 – 0.516 | 0.476 – 0.500 |
| Peron | 0.072 | - | 0.491 – 0.515 | 0.476 – 0.499 |
| Indulkanna | 0.486 | 0.489 | - | 0.013 – 0.017 |
| Raglessi | 0.477 | 0.481 | 0.015 | - |

Subsequent STRUCTURE analyses carried out within each species separately indicated there were two distinct clusters within the WGW, representing each of the populations that were sampled. Delta K and probability estimates also supported two clusters within the WGW (Figure 4). A small amount of admixture was observed in the Hamelin population cluster with some individuals showing shared ancestry with the Peron population cluster. The separation of Peron and Hamelin populations was also evident when individuals were assigned to three genetic clusters. Similar to the WGW, two genetic clusters were found that represented the two TBGW subspecies. Estimates of Delta K for both the TBGW and WGW analyses suggested K is two within each species (Figure 4). The STRUCTURE plot for the TBGW analysis where K = 2 showed that Indulkanna and Raglessi both had individuals with shared ancestry, suggesting some genetic admixture between these two TBGW subspecies (Figure 4). The results from the sensitivity analysis using conservative filtering parameters found that patterns of genetic structuring were consistent across all analyses (see Appendix S1).

**Figure 4.**
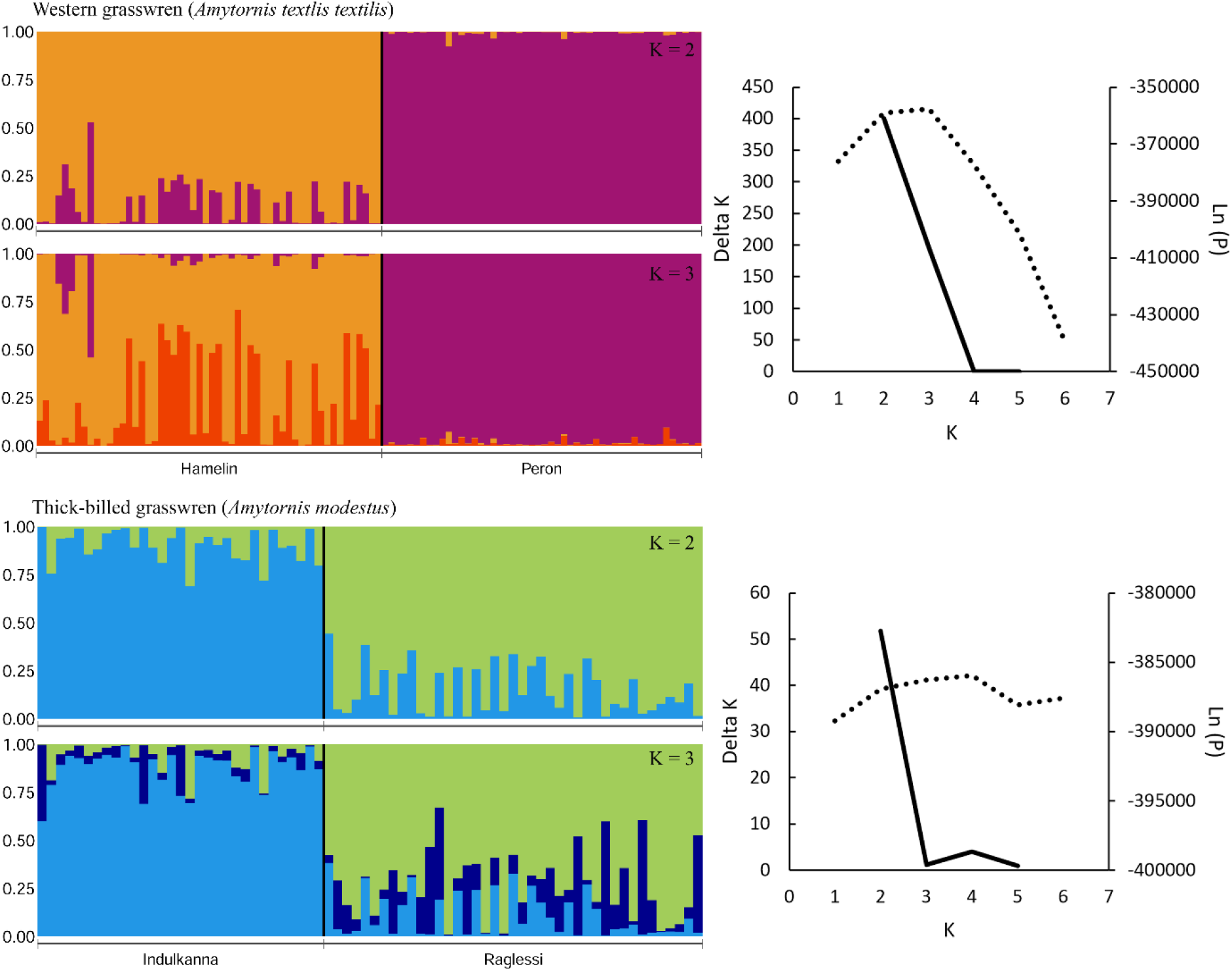
Summary of independent STRUCTURE analysis of the western grasswren (WGW; *Amytornis textilis textilis;* Hamelin and Peron) and thick-billed grasswren (TBGW; *Amytornis modestus;* Indulkanna and Raglessi), with genetic clusters represented by different colours. Each column represents an individual’s estimated allocation to each cluster. Accompanying each STRUCTURE plot are Delta K estimates for varying values of genetic clusters (K; solid line) and probability estimates for each K (Ln (P); dotted line) derived from the STRUCTURE analysis of WGW and TBGW.

### Spatial Autocorrelation Analysis

Evidence of genetic structure was observed at various distance classes in both grasswren species (Figure 5). Significantly positive values of *r* were found within both WGW populations (Peron and Hamelin) at the 0-5 km distance class, as well as the 5-15 km and 20-25 km distance classes for Hamelin (Figure 5). Similarly, significantly positive *r* values were found within both TBGW populations (Indulkanna and Raglessi) at the 0-5 km distance class and the 5-10 km and > 55 km distance classes in the Indulkanna samples (Figure 5).

**Figure 5.**
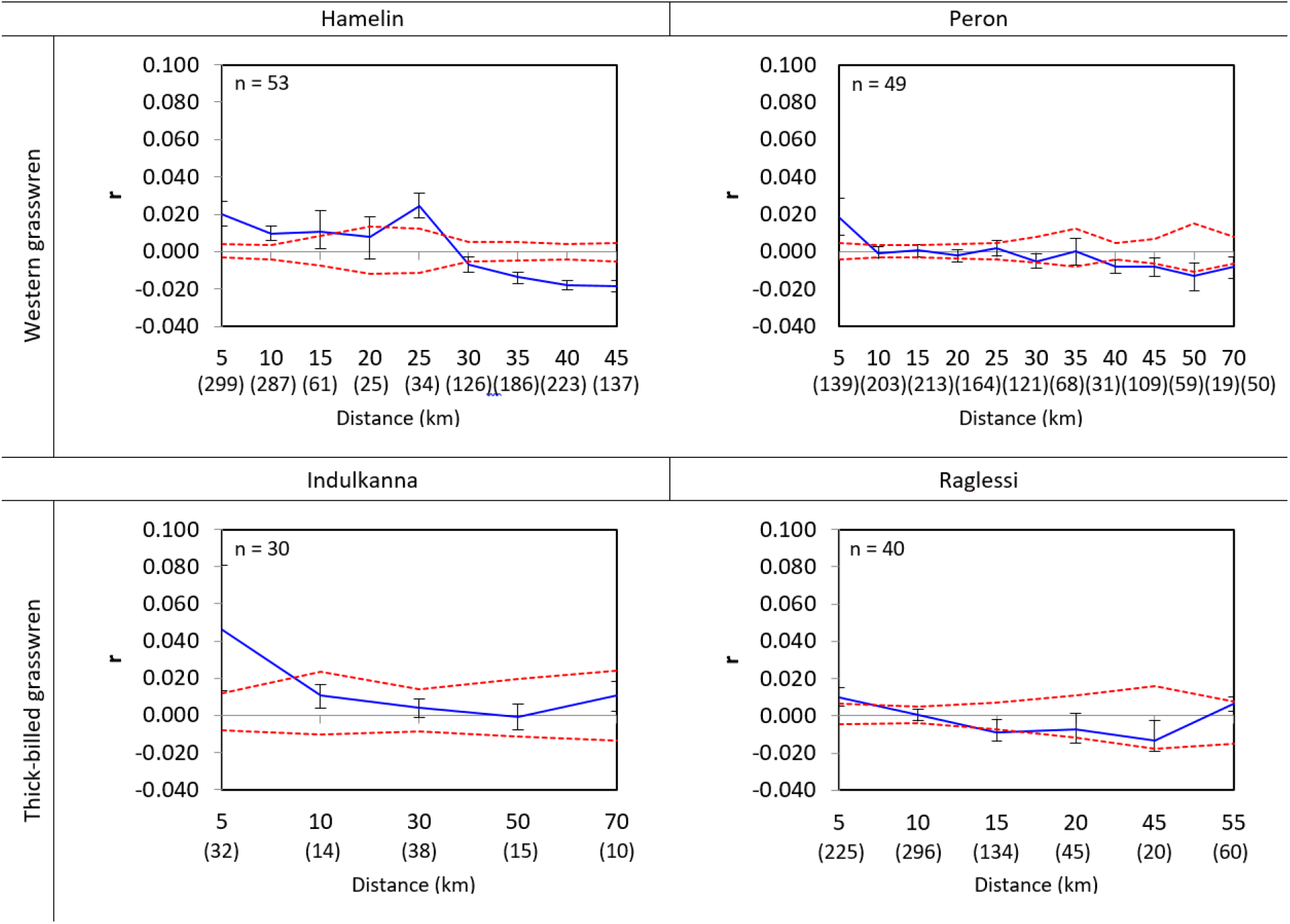
Spatial genetic autocorrelation (*r*) at variable distance classes for four populations of two grasswren species: western grasswren *(Amytornis textilis textilis;* Hamelin and Peron) and thick-billed grasswren *(Amytornis modestus;* Indulkanna and Raglessi). Dotted lines represent upper and lower 95% confidence intervals about the null hypothesis of no genetic structure. Error bars bound the 95% confidence interval about *r* as determined by bootstrap sampling. Pairwise comparison sample sizes for each distance class are in parentheses. Distance classes are in kilometres, and are variable between each plot. Each distance class is designated by the upper bound of the range it covered.

Examination of finer scale spatial autocorrelation within WGW revealed significantly positive *r* values between 0 and 500 m in both sexes in the Hamelin and Peron populations. In addition, significant positive *r* values were observed in females at Hamelin in distances classes greater than 10 km, and for males at the 10 km distance class, but this pattern was not observed in the Peron population (Figure 6).

**Figure 6.**
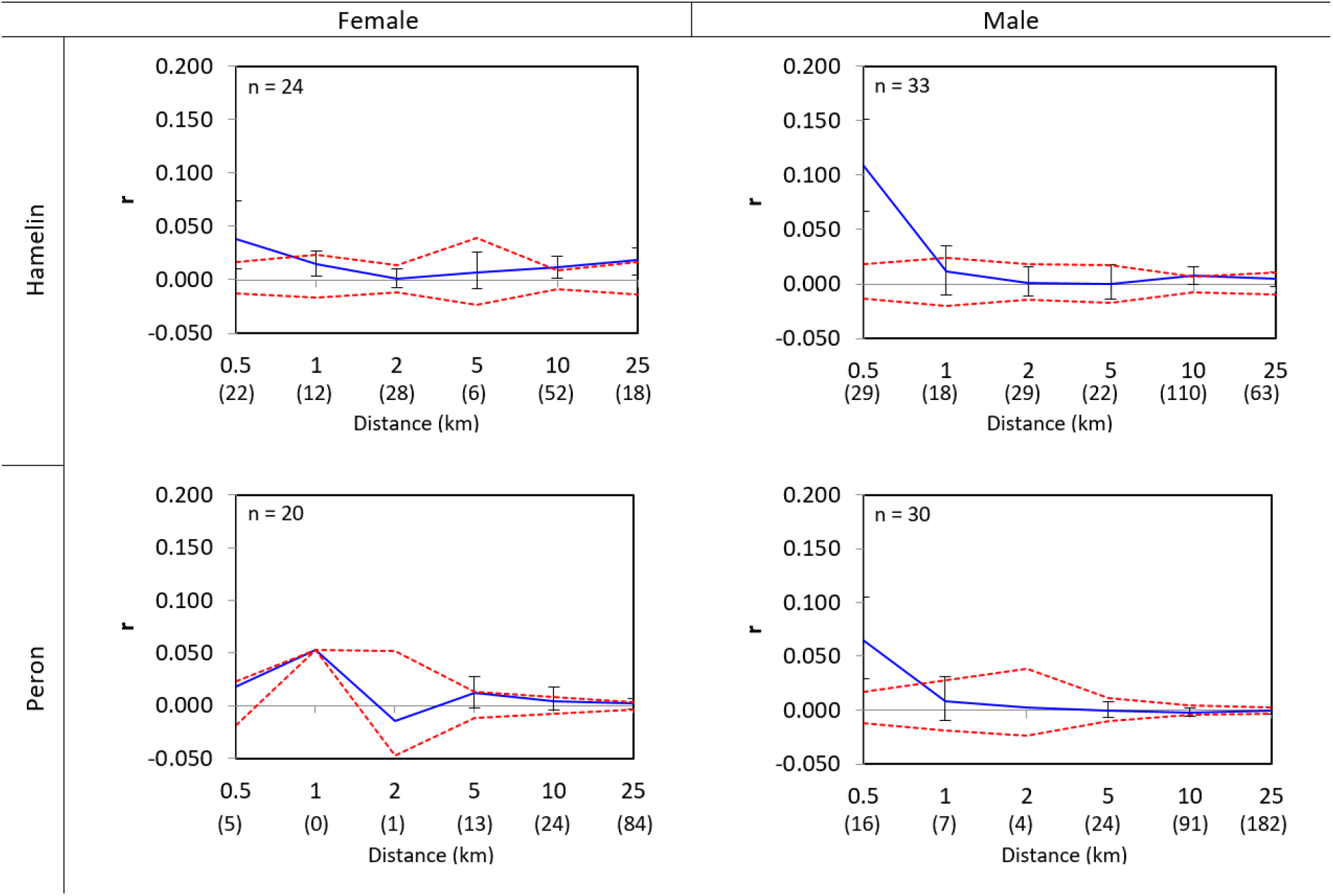
Spatial genetic autocorrelation at variable distance classes for the western grasswren *(Amytornis textilis textilis)* at two populations (Hamelin and Peron) for each sex. Dotted line represents upper and lower 95% confidence intervals about the null hypothesis of no genetic structure. Error bars bound the 95% confidence interval about *r* as determined by bootstrap sampling. Sample sizes for each distance class are in parentheses. Distance classes are in kilometres. Each distance class is the upper bound range at that given point.

### Morphological differences

There were significant differences between populations and sexes for all morphological traits across WGW and TBGW (see Appendix S3 for full table). For body mass, there were significant effects of population, sex and their interaction (*X*^2^ = 259.1, *p* = < 0.001, df = 3; *X*^2^ = 33.6, *p* = < 0.001, df = 1; *X*^2^ = 8.09, *p* = < 0.05, df = 3, respectively; Appendix S3). For all measured traits, male grasswrens were larger than female grasswrens of the same population (see Appendix S3). No significant intraspecific morphological differences were found for body mass and wing length in either species, but Peron birds had significantly larger head-bill lengths than Hamelin birds in both sexes (females, p < 0.01; males, p < 0.05; Appendix S3). Male Raglessi had longer tarsus length than Indulkanna (p < 0.01; Appendix S3), while the WGW populations were not morphologically divergent in this trait (p > 0.05). Raglessi females had significantly longer tails than Indulkanna (p < 0.01; Appendix S3).

The magnitude of interpopulation morphological differences within the two grasswren species varied among morphological traits (Table 4). WGW had a greater magnitude of divergence between the two populations than the TBGW for head-bill (both sexes) and tail length (males). In all other traits, the TBGW had greater morphological divergence between Indulkanna and Raglessi than the WGW populations (Table 4). However, the confidence intervals were very large and thus no definitive conclusions could be made about the variation in *P*_ST_ values between WGW and TBGW.

**Table 4.** Pairwise P_ST_ between populations for the western grasswren (WGW; Hamelin and Peron) and the thick-billed grasswren (TBGW; Indulkanna and Raglessi) for various morphological traits. Bootstrapped 95% confidence intervals are given in parenthesis.

| Trait | Female |  | Male |  |
| --- | --- | --- | --- | --- |
|  | WGW | TBGW | WGW | TBGW |
| Weight | 0.463 (0.005 – 0.836) | 0.764 (0.152 – 0.925) | 0.005 (0.0004 – 0.721) | 0.053 (0.001 – 0.738) |
| Head-bill | 0.735 (0.111 – 0.950) | 0.513 (0.003 – 0.871) | 0.857 (0.519 – 0.942) | 0.637 (0.014 – 0.885) |
| Tail length | 0.699 (0.075 – 0.912) | 0.750 (0.215 – 0.911) | 0.755 (0.059 – 0.940) | 0.626 (0.022 – 0.888) |
| Tarsus length | 0.717 (0.066 – 0.913) | 0.842 (0.463 – 0.951) | 0.211 (0.0004 – 0.765) | 0.855 (0.633 – 0.935) |
| Wing length | 0.463 (0.075 – 0.912) | 0.947 (0.897 – 0.971) | 0.709 (0.081 – 0.902) | 0.798 (0.349 – 0.922) |

## Discussion

We found strong genetic structuring within both grasswren species. Our finding of genetic divergence between the TBGW subspecies is consistent with a previous SNP study focussing on this species only (Slender *et al*. 2021). However, genetic structuring between the two WGW populations was much greater than between the TBGW subspecies, despite the WGW occurring over a smaller geographical area (150 km diameter) and having a smaller sampling gap (50 km) between the intraspecific populations. The genetic distinctiveness between the Peron and Hamelin WGW populations aligns with the genetic divergences found between Peron Peninsula and the Shark Bay mainland in the White-winged Fairywren (Walsh *et al*. 2021), supporting the hypothesis that peninsula populations are genetically isolated from adjacent mainland populations. Knowing that the two grasswren species have different evolutionary histories (Austin *et al*. 2013; Slender *et al*. 2021) further supports the hypothesis as the WGW subspecies is considered to be one ESU, and yet its populations are just as divergent, if not more, than that of what is considered to be two ESUs in the TBGW. This comparison highlights the impact of peninsulas on gene flow, and the potential for Peron and Hamelin to benefit from conservation management as independent ESUs. Our study also confirmed the genetic distinctiveness between WGW and TBGW, consistent with previous mitochondrial DNA studies based on limited historical samples (Austin *et al*. 2013; Christidis *et al*. 2013).

Genetic structuring within the WGW aligns closely with a geographical barrier that likely restricts gene flow. Peron is connected to Hamelin by a narrow 2.5 km wide corridor of land. This may greatly restrict dispersal between the Peron and Hamelin populations as grasswrens are not expected to be able to traverse water due to their wing morphology and have a tendency to move across the ground rather than fly (Rayner 1988). While previous records do show WGW south of the isthmus (Brooker 2000), the narrow corridor of land would still restrict gene flow as it creates a bottleneck dispersal effect limiting the amount of gene flow between populations. Isolation may also be impacting genetic diversity within the two WGW populations, which were significantly lower than the levels observed in Raglessi and Indulkanna. In contrast, the TBGW occupies a habitat that occurs heterogeneously over the landscape with intermittent dune fields that may restrict gene flow (Slender *et al*. 2017b). TBGW are also not constrained by coastline (i.e. open water between habitats), which may be a greater dispersal barrier to grasswrens as seen in the WGW. It is also known that the TBGW subspecies are in geographic contact (Slender 2018). All grasswrens are thought to be poor dispersers as they are small, mostly ground-dwelling (Christidis *et al*. 2010) and their short round wings likely limit long distance flight (Rayner 1988). Thus, our results on the WGW may be reflective of innately poor dispersal tendencies. To tease out the effects of geographic barriers versus inherent dispersal capabilities, this study incorporated consideration of spatial autocorrelation.

Our spatial autocorrelation analyses suggest male WGW display higher levels of philopatry than females, which is consistent with in-field observations of dispersed WGW males found near their natal territories (observations by A. Gibson Vega). High levels of philopatry observed in WGW males aligns with previous studies of closely related species within the family Maluridae (e.g. fairywrens; Cockburn et al., 2003; Double et al., 2005; Leitão et al., 2019). Our dataset indicates a majority of WGW males disperse short distance (less than 500 m). In contrast to males, female WGW appear to disperse longer distances from their natal territories (up to 25 km), consistent with the general trend in birds for females to disperse further from their natal area than males (Greenwood 1980). Habitat heterogeneity may explain the different spatial autocorrelation profiles we observed amongst the WGW populations. Greater habitat heterogeneity at Hamelin may be driving females to disperse further to find suitable territories and mates as they must cover a greater amount of unsuitable habitat between suitable patches compared to females from Peron. A second explanation could be that dispersal of Hamelin birds is not limited by obvious landscape features, while Peron birds are bound by coastline, and therefore have dispersal limitations. Our results, coupled with the resighting of a Hamelin colour-banded female grasswren approximately 30 km from the nearest banded bird at Hamelin confirms that WGW females can move distances of tens of kilometres (R. McLellan, personal communication, 2020). However, the STRUCTURE results show low levels of mixed ancestry in some Hamelin birds, suggesting that gene low can occur over larger distances and that Peron is the source of most gene flow between the two WGW populations. Those individuals showing highest mixed ancestry were evenly sampled through the geographic extent of the sampled Hamelin population. The even sampling provides evidence of infrequent, but present, directional dispersal between the entirety of Hamelin and Peron. Our spatial autocorrelation analyses on the TBGW populations were also indicative of long-distance dispersal, as positive spatial autocorrelation was detected beyond 55 km. These patterns of dispersal are consistent with results observed in a larger dataset of TBGW samples using the same methods (Slender *et al*. 2021). Our dataset and that of Slender et al. (2021) therefore suggest that grasswrens, despite being small ground-dwelling birds, are capable of dispersal over tens of kilometres in search of suitable habitat and mates. These results are consistent with the notion that genetic divergence between Hamelin and Peron populations is not due to poor dispersal capabilities, but rather to geographic landforms restricting dispersal.

Restricted dispersal between populations and the historical decline in the number of WGW may be the driver for the low genetic diversity found in the species (Frankham 1996). TBGW have a larger population and distribution with greater levels of connectivity, which may have contributed to higher levels of genetic variation in this species. However, interestingly, the TBGW had a greater deficiency of heterozygotes (significantly positive *F*_IS_) than the WGW despite having greater genetic diversity. This pattern may not be a true reflection of inbreeding, as *F*_IS_ is often interpreted to be, but rather a reflection of our sampling design (Wang 2014). Due to the complexities of capturing grasswrens, many samples often were collected alongside neighbours, which may have contributed to a deficiency in heterozygotes due to philopatry. Therefore, while we found genetic divergence and low genetic diversity between WGW populations, levels of inbreeding within the population remains inconclusive. Further investigation of sampled breeding pairs would inform whether inbreeding is actively occurring in WGW populations.

In addition to genetic divergence, morphological divergence was found in WGW and TBGW. Morphological divergence has been previously described in the TBGW, and was a contributing factor to the current subspecies classification (Black 2011a; Slender *et al*. 2017a). Morphological divergences between the two WGW populations were similar in magnitude to those found between the TBGW populations for some traits (e.g. male wing length and female tail length). However, these results should be interpreted cautiously given the large confidence intervals for most *P*_ST_ estimates. For example, our results indicate that Raglessi males have longer tarsi than Indulkanna males, and that Raglessi females have longer tails than males, which contradicts previous work done by Slender et al. (2017a) on morphology, using a larger sample size. This may once again reflect that our morphological data was not robust and results should be interpreted cautiously. As such, it is unclear whether the degree of morphological divergence was similar across species, and whether it reflected the same magnitude of genetic divergence.

Understanding genetic structure (and hence landscape barriers), genetic diversity, and dispersal can assist in identifying populations of conservation priority (e.g. von Takach *et al*., 2021). The WGW requires such relevant ecological understanding for the development of conservation and management strategies. A translocation is planned for this species to an island near Shark Bay (Algar *et al*. 2020). WGW subspecies *Amytornis textilis carteri* once occurred on Dirk Hartog Island (also in Shark Bay), but is believed to have gone extinct due to land degradation from overgrazing by introduced stock and goats, and possibly due to predation by feral cats (Cale 2003; Black *et al*. 2021). There are plans to utilise the mainland WGW subspecies as an ecological replacement as part of a wider reconstruction of the island’s faunal assemblage (Algar *et al*. 2020), but data on most aspects of Western Grasswren ecology are currently lacking to develop an informed translocation plan. The IUCN/SSC *Guidelines for Reintroductions and Other Conservation Translocations* (IUCN/SCC 2013) outline genetic considerations for conservation translocations, including aiming to provide adequate genetic diversity, mixing populations to maximise diversity, while minimising the risk of genetic incompatibilities. The planned WGW translocation may benefit from sourcing from both populations (Hamelin and Peron) to increase genetic diversity and retain adaptive potential of the founding population on Dirk Hartog Island (Binks *et al*. 2007; Kennington *et al*. 2012; Thavornkanlapachai *et al*. 2019b). The latter is of particular importance as what is deemed suitable habitat for WGW on Dirk Hartog Island is not identical to habitat present in Peron or Hamelin (Brooker 2000; Black 2011b; A. Gibson Vega, personal observation)

## Conclusion

Our study has shown that the genetic divergence between two WGW populations in close proximity exceeds the level found between two described subspecies of TBGW, which are genetically and morphologically divergent. Comparison of WGW with the TBGW populations indicates that grasswrens can and do move relatively large distances, but may be constrained by coastlines such as those found in Shark Bay. Our results suggest that grasswrens are able to traverse unsuitable terrestrial habitat (as seen in the TBGW), but not open water (as seen in the WGW), which is to be expected as grasswrens are not morphologically adapted for long distance flight. The limited, but directional, gene flow between the two WGW populations further suggests that dispersal from Peron peninsula to Hamelin is more likely as Peron birds need to go through a bottle neck of land towards Hamelin. In contrast Hamelin birds have dispersal opportunities to move north, east or south, as well was towards Peron. Hence it’s more likely that birds from Peron and up at Hamelin than vice versa. Low genetic diversity within Peron and Hamelin highlights the importance of maintaining and restoring habitat in both parts of the overall distribution within WGW. Shark Bay is the only location for WGW in Western Australia, and while there has been no recent observed population decline in the Shark Bay area itself, the distribution of the remaining population of WGW in Western Australia is geographically limited and thus vulnerable to stochastic events. From a management perspective, our results show a need to place equal importance on the two WGW populations, as they are genetically distinct with little contemporary gene flow between them.

## Ethical approval

Samples were collected from Western Australia under Department of Biodiversity, Conservation and Attractions (DBCA) license FO25000186 (A. Burbidge). Animal ethics approval was granted by the DBCA (2019-05A) for Western Grasswren blood sample collection. Thick-billed Grasswren samples were collected under animal ethics approval from Flinders University Animal Welfare Committee (animal ethics approval E385). No animal was removed from its natural habitat for the purpose of this study.

## Author contributions

A.G.V was responsible for data collection of western grasswren samples, conducting the research, analysing data and the main contributor to the manuscript. W.J.K provided extensive revisions of the analysis and advised on the interpretation of the results. A.L.S and M.L. collected the thick-billed samples and provided revisions on the manuscript. M.L.H., A.R., S.J.C. and A.H.B. provided revisions to the entire text.

## Data archiving

SNP data is available upon request from the author

## Acknowledgements

We acknowledge the Malgana and Whadjuk Noongar people as the Traditional Owners of the Land on which this research was conducted. We are grateful to Bush Heritage Australia for permitting work on Hamelin Station Reserve, and providing valuable in-kind support and collaboration on this study. Particular thanks go to M. and K. Judd at Hamelin Station Reserve for their technical support at Hamelin Station Reserve, and B. Parkhurst for sharing knowledge of grasswren locations. We are thankful for the field assistance provided by C. Greenwell, C. Bowry, J. Ringma, I. Pereda, R. de Visser, G. Blackburn, K. Rayner, M. Blythman, L. Dadour, B. Parkhurst, G. Ricci and B. Richards. We are grateful to Y. Hitchen for her guidance in the lab. This research was supported by the University of Western Australia, Department of Biodiversity, Conservation and Attractions, Bush Heritage Australia and the Australian Bird and Bat Banding Scheme. This research was generously supported by the Holsworth Wildlife Endowment, Wettenhall Environmental Trust and Paul Hackett Memorial project grants, alongside valuable funding from the Gorgon Barrow Island Net Conservation Benefits Fund. A. Gibson Vega is supported by an Australian Government Research Training Program scholarship through the University of Western Australia.

