## Supplemental material for "A comparative study of two grasswren species reveals strong genetic divergence between a peninsula and mainland population"

**Appendix**

**Appendix S1**

**Table S1.1** Alternative conservative SNP filtering pipelines, used as a sensitivity analysis to demonstrating that patterns of genetic structuring was consistent across all analyses.

| Parameters | *Conservative* | *Very conservative* |
| --- | --- | --- |
| Call rate | 0.9 | 0.85 |
| Minor allele frequency | 0.05 | 0.02 |
| Monomorphs | Excluded | Excluded |
| Outliers identified and excluded | 15 | 17 |
| Loci remaining | 11 282 | 21 479 |

**Table S1.2**. Observed and expected heterozygosity, and inbreeding coefficient (F_IS_) for *conservative* and *very conservative* SNP filtering pipelines. Asterisks indicate significant deviation from zero. While the raw values are different for the two filtering pipelines, the patterns between WGW (Hamelin and Peron) and TBGW (Indulkanna and Raglessi) remain the same.

|  |  | *Conservative* | | | *Very conservative* | | | |
| --- | --- | --- | --- | --- | --- | --- | --- | --- |
| Population | N | Ho (± SE) | He (± SE) | F_IS_ (± SE) | N | Ho (± SE) | He (± SE) | F_IS_ (± SE) |
| Hamelin | 52 | 0.115 ± 0.001 | 0.141 ± 0.002 | 0.155 ± 0.002* | 52 | 0.085 ± 0.001 | 0.107 ± 0.001 | 0.164 ± 0.002* |
| Peron | 49 | 0.122 ± 0.001 | 0.145 ± 0.002 | 0.130 ± 0.002* | 49 | 0.088 ± 0.001 | 0.108 ± 0.001 | 0.141 ± 0.002* |
| Indulkanna | 31 | 0.172 ± 0.001 | 0.243 ± 0.002 | 0.265 ± 0.003* | 31 | 0.122 ± 0.001 | 0.186 ± 0.001 | 0.300 ± 0.002* |
| Raglessi | 41 | 0.172 ± 0.001 | 0.247 ± 0.002 | 0.278 ± 0.003* | 41 | 0.124 ± 0.001 | 0.191 ± 0.001 | 0.312 ± 0.002* |

A Friedman’s test revealed that there was a significant effect in observed and expected heterozygosity between populations in the *conservative* (H_O_; *X*^2^ = 1202.7, df = 3, p-value < 0.001, H_e_; *X*^2^ = 1720.4, df = 3, p-value < 0.001) and *very conservative* (H_O_; *X*^2^ = 1932.7, df = 3, p-value < 0.001, H_e_; *X*^2^ = 2985.5, df = 3, p-value < 0.001 ) datasets.

**Table S1**.**3** Pairwise Wilcoxon rank sum test results comparing western grasswren (*Amytornis textilis textilis;* Hamelin and Peron) and thick-billed grasswren (*Amytornis modestus;* Indulkanna, and Raglessi) populations for *conservative* and *very conservative* SNP filtering pipelines. Above the diagonal are p values comparing observed heterozygosity between populations, while below the diagonal p values are for expected heterozygosity.

|  |  | Hamelin | Peron | Indulkanna | Raglessi |
| --- | --- | --- | --- | --- | --- |
| *Conservative* | Hamelin | - | 0.24 | < 0.001 | < 0.001 |
|  | Peron | 0.68 | - | < 0.001 | < 0.001 |
|  | Indulkanna | < 0.001 | < 0.001 | - | 0.49 |
|  | Raglessi | < 0.001 | < 0.001 | 0.64 | - |
| *Very conservative* |  |  |  |  |  |
|  | Hamelin | - | 0.22 | < 0.001 | < 0.001 |
|  | Peron | 0.006 | - | < 0.001 | < 0.001 |
|  | Indulkanna | < 0.001 | < 0.001 | - | < 0.001 |
|  | Raglessi | < 0.001 | < 0.001 | < 0.001 | - |

**Table S1.4**. Summary of pairwise FST results for the various sampling populations for the western grasswren (*Amytornis textilis*; Hamelin and Peron) and thick-billed grasswren (*Amytornis modestus*; Indulkanna, Raglessi and Hybrid Zone) for *conservative* and *very conservative* datasets. Below the diagonal are the Nei FST results. Above the diagonal are 95% confidence interval bootstrap results.

|  |  | Hamelin | Peron | Indulkanna | Raglessi |
| --- | --- | --- | --- | --- | --- |
| *Conservative* | Hamelin | - | 0.070 – 0.077 | 0.497 – 0.516 | 0.481 – 0.499 |
|  | Peron | 0.073 | - | 0.498 – 0.518 | 0.483 – 0.502 |
|  | Indulkanna | 0.488 | 0.492 | - | 0.014 – 0.017 |
|  | Raglessi | 0.481 | 0.485 | 0.16 | - |
| *Very conservative* | Hamelin | - | 0.067 – 0.074 | 0.463 – 0.481 | 0.445 – 0.462 |
|  | Peron | 0.071 | - | 0.467 – 0.484 | 0.449 – 0.466 |
|  | Indulkanna | 0.453 | 0.458 | - | 0.015 – 0.017 |
|  | Raglessi | 0.443 | 0.448 | 0.016 | - |

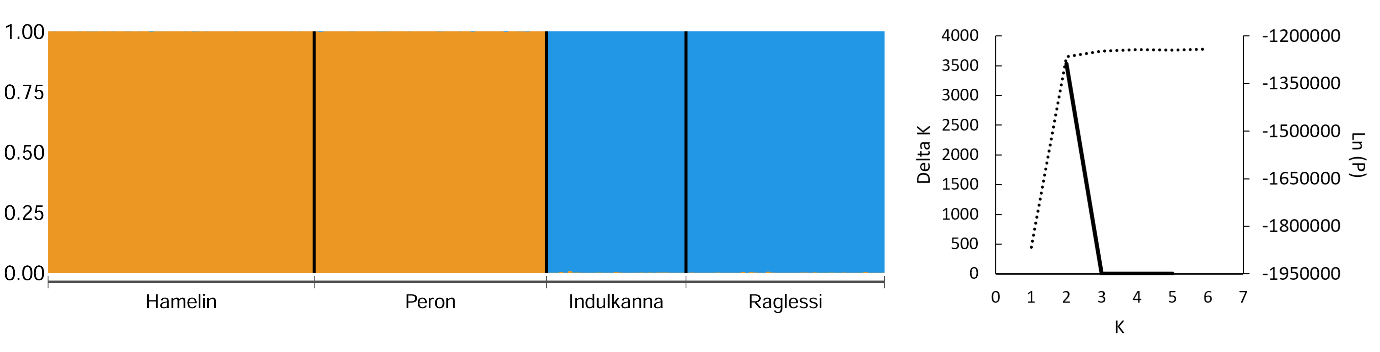

**Figure S1.5.** Summary of the STRUCTURE analysis utilising the *conservative* dataset of the western grasswren (*Amytornis textilis textilis*; Hamelin and Peron) and thick-billed grasswren (*Amytornis modestus*; Indulkanna, Raglessi and Hybrid Zone), with genetic clusters represented by different colours (left). Δ K estimates for varying values of genetic clusters (K; solid line) and probably estimates for each K (Ln (P); dotted line), derived from the STRUCTURE analysis.

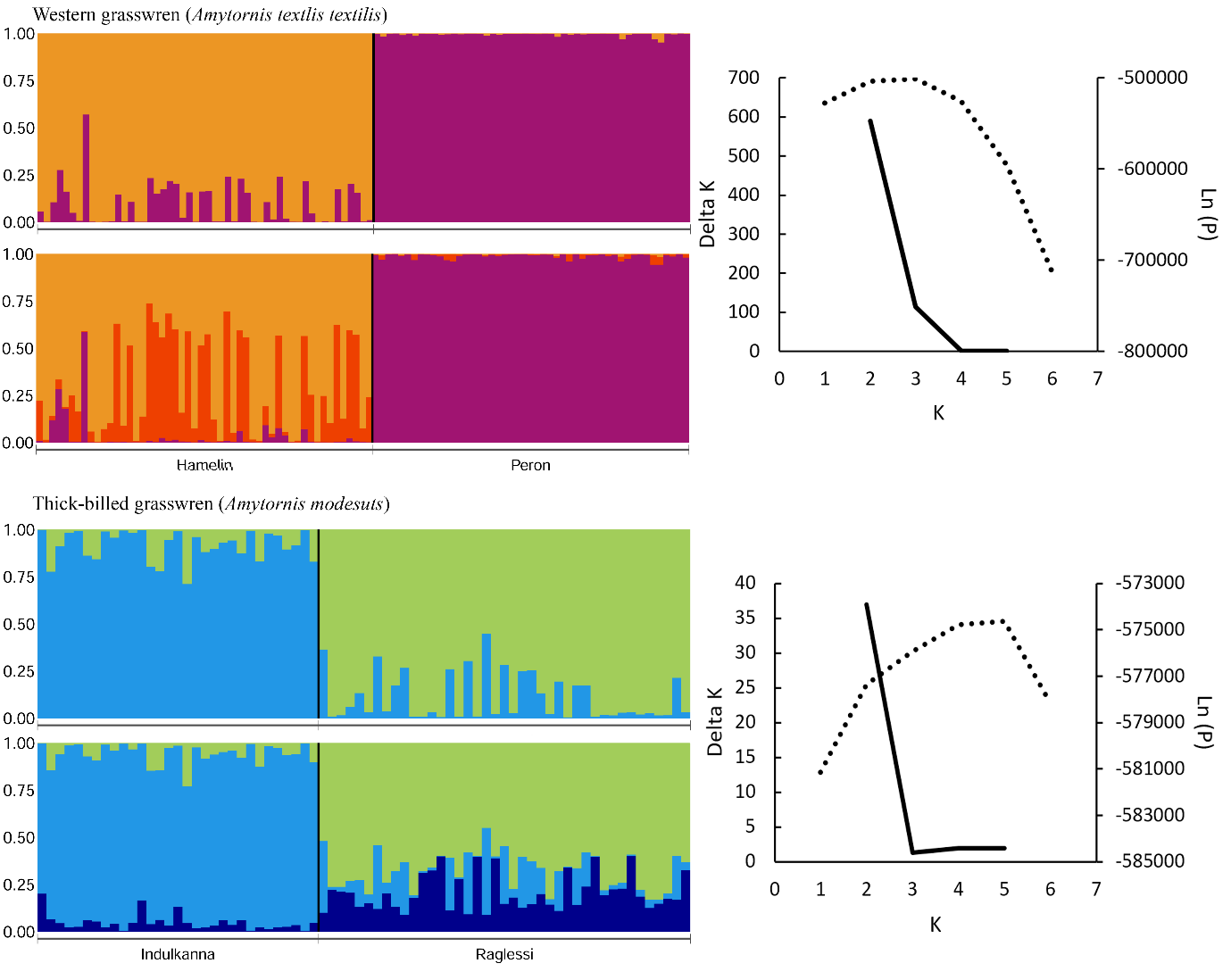

**Figure S1.6**. Summary of independent STRUCTRE analysis utilising the *conservative* dataset of the western grasswren (WGW; *Amytornis textilis textilis*; Hamelin and Peron) and thick-billed grasswren (TBGW; *Amytornis modestus*; Induklanna, Raglessi and Hybrid Zone), with genetic clusters represented by different colours. The top plot is K = 2 and the bottom represents K = 3, for each species. Accompanying each STRUCTURE plot are Δ K estimates for varying values of genetic clusters (K; solid line) and probability estimates for each K (Ln (P); dotted line) derived from the STRUCTURE analysis of WGW and TBGW.

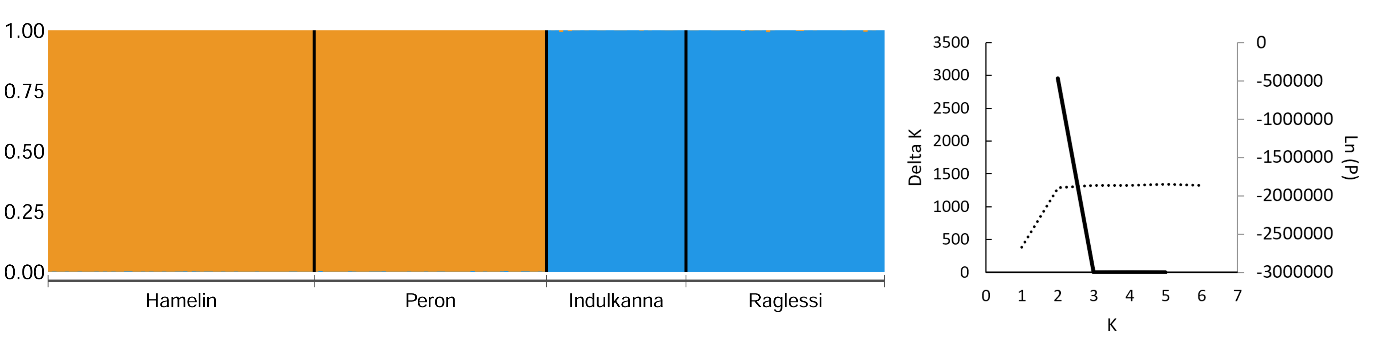

**Figure S1.7.** Summary of the STRUCTURE analysis utilising the very *conservative* dataset of the western grasswren (*Amytornis textilis textilis*; Hamelin and Peron) and thick-billed grasswren (*Amytornis modestus*; Indulkanna, Raglessi and Hybrid Zone), with genetic clusters represented by different colours (left). Δ K estimates for varying values of genetic clusters (K; solid line) and probably estimates for each K (Ln (P); dotted line), derived from the STRUCTURE analysis.

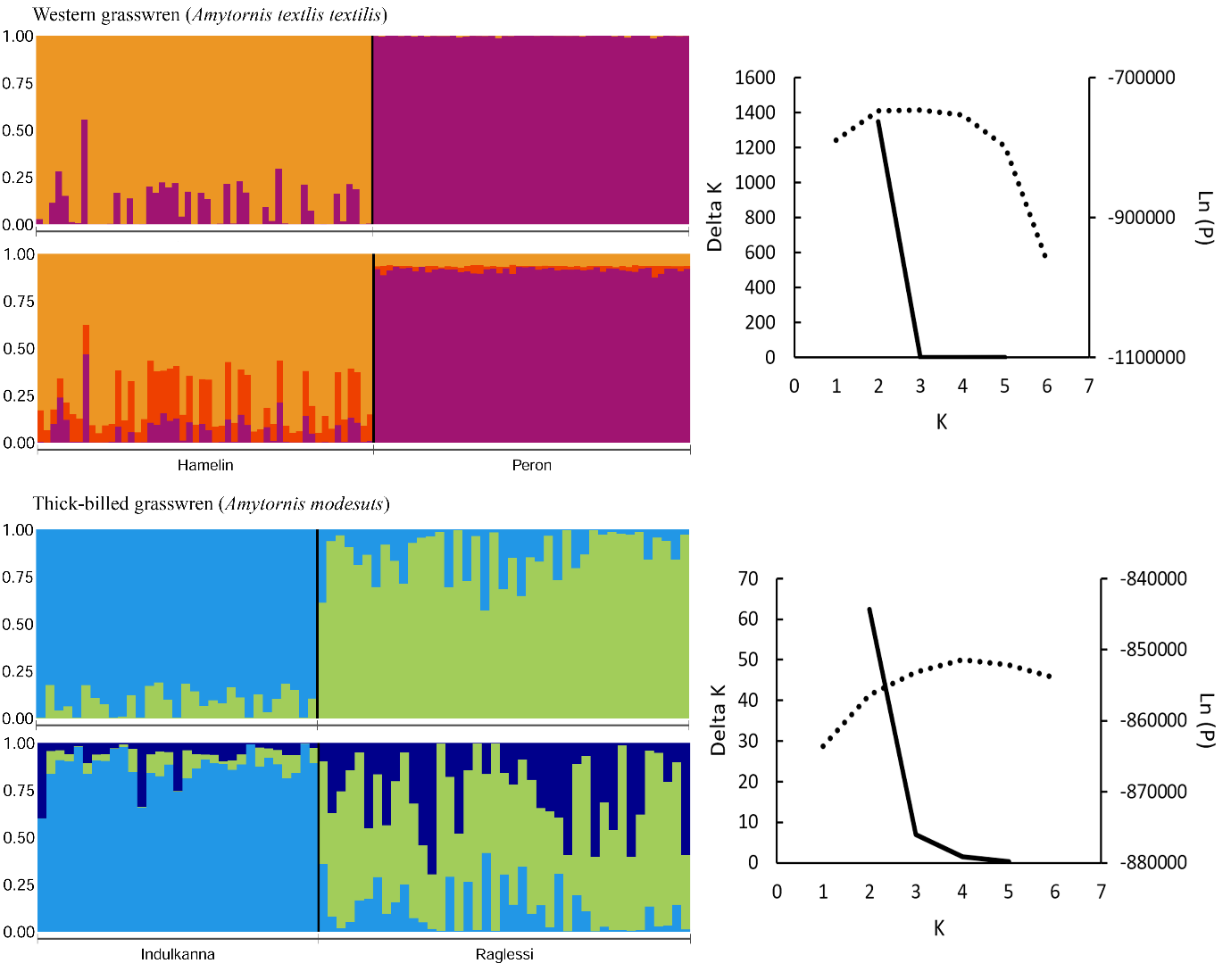

**Figure S1.8**. Summary of independent STRUCTRE analysis utilising the *very* *conservative* dataset of the western grasswren (WGW; *Amytornis textilis textilis*; Hamelin and Peron) and thick-billed grasswren (TBGW; *Amytornis modestus*; Induklanna, Raglessi and Hybrid Zone), with genetic clusters represented by different colours. The top plot is K = 2 and the bottom represents K = 3, for each species. Accompanying each STRUCTURE plot are Δ K estimates for varying values of genetic clusters (K; solid line) and probability estimates for each K (Ln (P); dotted line) derived from the STRUCTURE analysis of WGW and TBGW.

**Appendix S2**

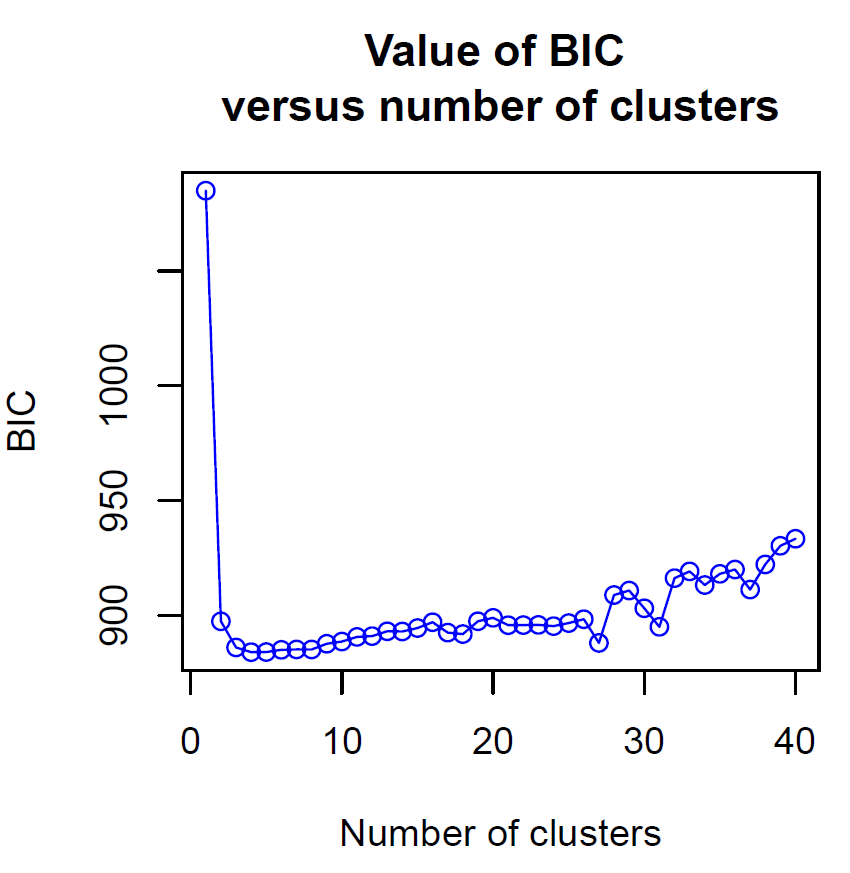

**Figure S2.1.**  BIC against different number of clusters for a DAPC analysis of western grasswren (*Amytornis textilis textilis*) and thick-billed grasswren (*Amytornis modestus*) populations. Optimal genetic clusters for this analysis is between 2 and 3 genetic clusters.  **Appendix S3**

**Table S3.1.** Effect of population (WGW; Peron and Hamelin. TBGW; Indulkanna and Raglessi), sex and their interaction on various morphological traits. Bold denotes statistical significance of p < 0.05.

| Trait | Effect | *X*^2^ | Df | p |
| --- | --- | --- | --- | --- |
| Body mass | Sex | 33.631 | 1 | **< 0.001** |
|  | Pop | 259.123 | 3 | **< 0.001** |
|  | Sex:Pop | 8.088 | 3 | **< 0.05** |
| Head-bill length | Sex | 50.903 | 1 | **< 0.001** |
|  | Pop | 65.623 | 3 | **< 0.001** |
|  | Sex:Pop | 5.83 | 3 | 0.122 |
| Tail length | Sex | 47.120 | 1 | **< 0.001** |
|  | Pop | 946.512 | 3 | **< 0.001** |
|  | Sex:Pop | 0.833 | 3 | 0.842 |
| Tarsus length | Sex | 44.245 | 1 | **< 0.001** |
|  | Pop | 254.870 | 3 | **< 0.001** |
|  | Sex:Pop | 5.917 | 3 | 0.116 |
| Wing length | Sex | 60.768 | 1 | **< 0.001** |
|  | Pop | 111.022 | 3 | **< 0.001** |
|  | Sex:Pop | 4.045 | 3 | 0.257 |

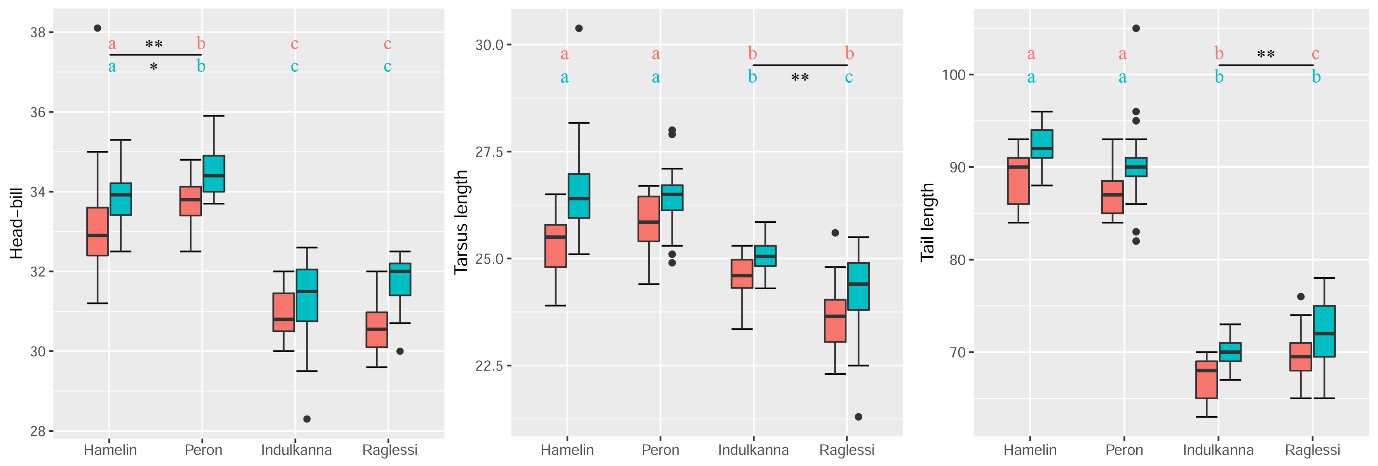

**Figure S3.2**. Morphological traits which have significant intraspecific population differences. Females are in pink, while males are in blue. Letters above correspond to significant differences between populations, with the colour of the text indicating sex. Intraspecific differences were determined by a Tukey’s Honest Significance Difference test, where asterisks denote significance (* = 0.05, ** = 0.01) morphological differences within the species (above the line is female significance, below the line is male significance).

**Table S3.3**. Means and standard error (in parenthesis) for various morphological traits in the western grasswren (Hamelin and Peron) and the thick-billed grasswren (Indulkanna and Raglessi).

| Trait |  | Hamelin | Peron | Indulkanna | Raglessi |
| --- | --- | --- | --- | --- | --- |
| Weight | Female | 24.114 (± 0.439) | 23.313 (±0.539) | 19.636 (±0.501) | 18.233 (±0.372) |
|  | Male | 25.588 (±0.284) | 25.548 (±0.362) | 19.716 (±0.257) | 19.592 (±0.293) |
| Head-bill | Female | 33.093 (±0.238) | 33.767 (±0.166) | 30.921 (±0.178) | 30.6 (±0.173) |
|  | Male | 33.911 (±0.122) | 34.473 (±0.131) | 31.237 (±0.255) | 31.736 (±0.147) |
| Tail length | Female | 88.879 (±0.496) | 87.210 (±0.680) | 67.077 (±0.582) | 69.341 (±0.769) |
|  | Male | 92.348 (±0.360) | 90.231 (±0.985) | 70.0 (±0.412) | 71.708 (±0.857) |
| Tarsus length | Female | 25.382 (±0.126) | 25.815 (±0.192) | 24.468 (±0.175) | 23.666 (±0.202) |
|  | Male | 26.548 (±0.174) | 26.385 (±0.159) | 25.074 (±0.102) | 24.212 (±0.234) |
| Wing length | Female | 64.466 (±0.343) | 63.333 (±0.443) | 56.385 (±0.279) | 59.312 (±0.434) |
|  | Male | 66.265 (±0.341) | 65.145 (±0.482) | 58.947 (±0.310) | 60.536 (±0.496) |
